# Perceptual similarity judgments do not predict the distribution of errors in working memory

**DOI:** 10.1101/2022.05.24.493200

**Authors:** Ivan Tomić, Paul M. Bays

## Abstract

Population coding models provide a quantitative account of visual working memory (VWM) retrieval errors with a plausible link to the response characteristics of sensory neurons. Recent work has provided an important new perspective linking population coding to variables of signal detection, including d-prime, and put forward a new hypothesis: that the distribution of recall errors on, e.g., a color wheel, is a consequence of the psychological similarity between points in that stimulus space, such that the exponential-like psychophysical distance scaling function can fulfil the role of population tuning and obviate the need to fit a tuning width parameter to recall data. Using four different visual feature spaces, we measured psychophysical similarity and memory errors in the same participants. Our results revealed strong evidence for a common source of variability affecting similarity judgments and recall estimates, but did not support any consistent relationship between psychophysical similarity functions and VWM errors. At the group level, the responsiveness functions obtained from the psychophysical similarity task diverged strongly from those that provided the best fit to working memory errors. At the individual level, we found convincing evidence against an association between observed and best-fitting similarity functions. Finally, our results show that the newly proposed exponential-like responsiveness function has in general no advantage over the canonical von Mises (circular normal) function assumed by previous population coding models.

## Perceptual similarity judgments do not predict the distribution of errors in working memory

When asked to reproduce simple visual features from memory, human observers give responses that deviate in a systematic manner from correct values. Efforts to understand the nature of these recall errors have led to the development of numerous quantitative models of VWM recall aimed at accounting for the shape of empirical error distributions, which have excess kurtosis compared to a normal distribution, with more density in the tails, an effect that becomes more prominent as the number of memoranda increases. Despite a long-running dispute about how and why these errors arise (Bays, 2014; Fougnie, Suchow, & Alvarez, 2012; Luck & Vogel, 1997; van den Berg, Shin, Chou, George, & Ma, 2012; Zhang & Luck, 2008), recent attempts to unify existing models within the framework of sampling have shown that dominant models of VWM share previously unrecognised commonalities (Schneegans, Taylor, & Bays, 2020). Here, we provide evidence for a further correspondence between influential models of VWM.

Computational theories of population coding (Pouget, Dayan, & Zemel, 2000, 2003) form the basis of one successful approach to explaining the data and mechanisms of VWM recall. In the Neural Resource model (Bays, 2014, 2015; Bays & Taylor, 2018; Schneegans & Bays, 2017), visual features are first encoded in and subsequently reconstructed from the activity of a population of idealized feature-selective neurons. In the simplest instantiation of the model (depicted in Fig. 1a–d), all neurons are assumed to have identical bell-shaped (von Mises) tuning curves of width *W*, translated through the feature space to peak at each neuron’s individual preferred feature value, such that they provide dense uniform coverage of the feature space (Fig. 1a). The joint response of the population to a particular stimulus can be visualized as a hill of activity with the same width and shape as the neural tuning curve, centered on the true stimulus feature and scaled by a peak amplitude (*r*_*max*_) (Fig. 1b). At read-out this mean response is translated into discrete spike counts with Poisson variability (Fig. 1c), which are decoded according to the principle of maximum likelihood to obtain a recall estimate.

**Figure 1.**
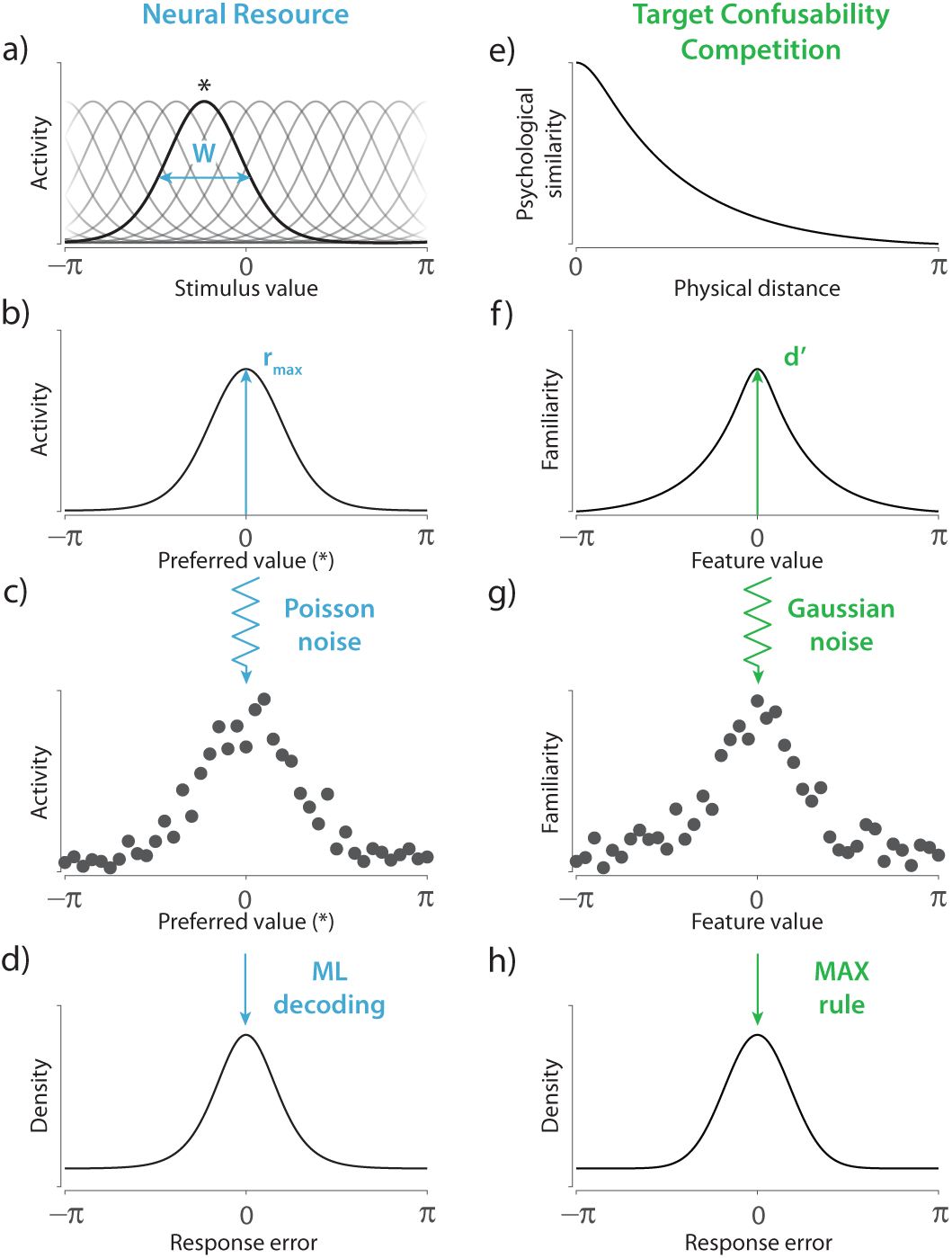
Comparison of the Neural Resource model and the Target Confusability Competition (TCC) model. *Note*. **(a)** In the Neural Resource model, stimuli are encoded in an idealized homogeneous population of spiking neurons whose activation depends on their individual tuning preferences and common tuning width (*W*). Asterisk indicates the preferred value of the neuron with the highlighted tuning function. **(b)** Activation as a function of preferred value, responding to a stimulus value of zero. The amplitude of the stimulus-evoked response is determined by the peak firing rate (*r*_*max*_). **(c)** Spikes are generated according to a Poisson process. **(d)** Recall estimates are obtained by a maximum likelihood decoder. The resulting distributions of error display long tails, consistent with patterns of human recall error. **(e)** In the TCC model, the psychological similarity between possible stimuli plays a critical role. **(f)** Recall of an item produces a pattern of familiarity signals that peaks over the encoded stimulus with strength *d*^*′*^, and spreads throughout the feature space in proportion to each values’ psychological similarity to the encoded stimulus. **(g)** The pattern of familiarity signals is corrupted by Gaussian noise. **(h)** The feature value with the strongest signal is reported. Like the population coding account, this model predicts long-tailed distributions of error.

The amplitude of activity controls the encoding precision, with larger values of *r*_*max*_ giving rise to more reliable decoded estimates. The width of the tuning curve is reflected in the distribution of reconstructed values, with narrower tuning producing stronger deviations from a normal distribution of error, including longer tails at low activity levels. A further assumption of the model is that the population activity encoding all stimuli is normalized, meaning that the total activity level is fixed and shared between memory items, implementing a form of flexible memory resource (Carandini & Heeger, 2011). The predictions of this model have proved a close fit to empirical human recall errors in analogue report tasks, and in particular quantitatively reproduce effects of set size and predictive cues on those errors.

A superficially quite different account of recall was recently proposed by Schurgin, Wixted, and Brady (2020), who argued that the shape of the WM error distribution can be explained by the psychological scaling of distances in the feature space. According to the Target Confusability Competition (TCC) model (depicted in Fig. 1e–h), the probability of confusing features in memory is determined by a function which maps distances between features on the physical scale to their perceived similarity (Fig. 1e). This function can be measured in a perceptual task in which observers make judgments about the relative similarity of pairs of stimuli (e.g., the method of quadruples; Maloney & Yang, 2003). As demonstrated by a large literature on similarity and generalization, these functions are approximately exponential with respect to physical distance (Nosofsky, 1992; Shepard, 1987).

In the TCC framework, recalling an item produces an internal response or familiarity signal with a peak amplitude (*d*^*′*^) at the true feature value, and a spread to neighbouring feature values that is proportional to the psychological similarity function (Fig. 1f). This mean response is corrupted by noise drawn from a standard normal distribution (Fig. 1g), and a recall estimate is obtained by applying a Max rule to the familiarity pattern, i.e., choosing the feature value with the strongest familiarity signal (Fig. 1h). Successful fits of this model have been demonstrated to error distributions in analogue report tasks, and also 2-AFC memory tasks.

Although described at different levels of analysis and using different concepts, there is a close correspondence between Neural Resource and TCC models. In both cases, the internal response to an encoded stimulus is a symmetrical function over possible feature values that peaks at the true stimulus value and decays with distance in stimulus space (Fig. 1b & f). And in both cases, a recall estimate is recovered from a noise-corrupted version of this ideal response function (Fig. 1c & g), such that recall precision is governed by the amplitude of the internal response (set by parameters *r*_*max*_ and *d*^*′*^, respectively).

This leaves the choice of responsiveness function as the substantive difference between accounts of WM, with TCC using a psychophysical similarity function, independently estimated in a perceptual task, in place of the Neural Resource model’s canonical von Mises function varying only in width. The seeming success of both models in replicating human memory errors raises the possibility of linking psychological similarity to principles of neural coding. However, the source study did not systematically examine whether variations in psychophysical similarity functions, either across individuals or feature spaces, corresponded to observed differences in the shape of WM error distributions. The goal of this study was to investigate and formally test the hypothesized relationship between judgments of perceptual similarity and working memory recall errors in a large group of participants and a range of visual feature spaces.

## Methods

### Participants

A total of 396 naive observers (205 females, 169 males; aged 18–35) took part in the study after giving informed consent in accordance with the Declaration of Helsinki. All observers were recruited using Prolific (https://www.prolific.co), reported normal color vision and normal or corrected-to-normal visual acuity, and were remunerated £5 per hour for their participation. Fifty-six observers initially recruited to the study were subsequently excluded and replaced with new participants for one of the following reasons: performance at chance levels (47), completing the task too fast (4), or technical errors (5). In total, 90 observers participated in the color study, 100 in the orientation study, 104 in the angular location study, and 102 in the shape study.

### General methods

Observers participated in three tasks: a working memory analogue report task, a perceptual analogue report task and a psychophysical scaling quad task. All observers completed tasks in the same order and on separate days. All observers completed the quad and WM task, and out of the total sample, 323 observers completed the perceptual analogue report task. The tasks were presented via browsers on observers’ personal computers and were coded in JavaScript and HTML Canvas.

### Stimuli

Four classes of stimuli were tested in separate studies: colors, orientations, angular locations, and artificial shapes, all drawn from respective continuous circular feature spaces (Figure 2). In all tasks, stimuli were presented against a mid-grey background.

**Figure 2.**
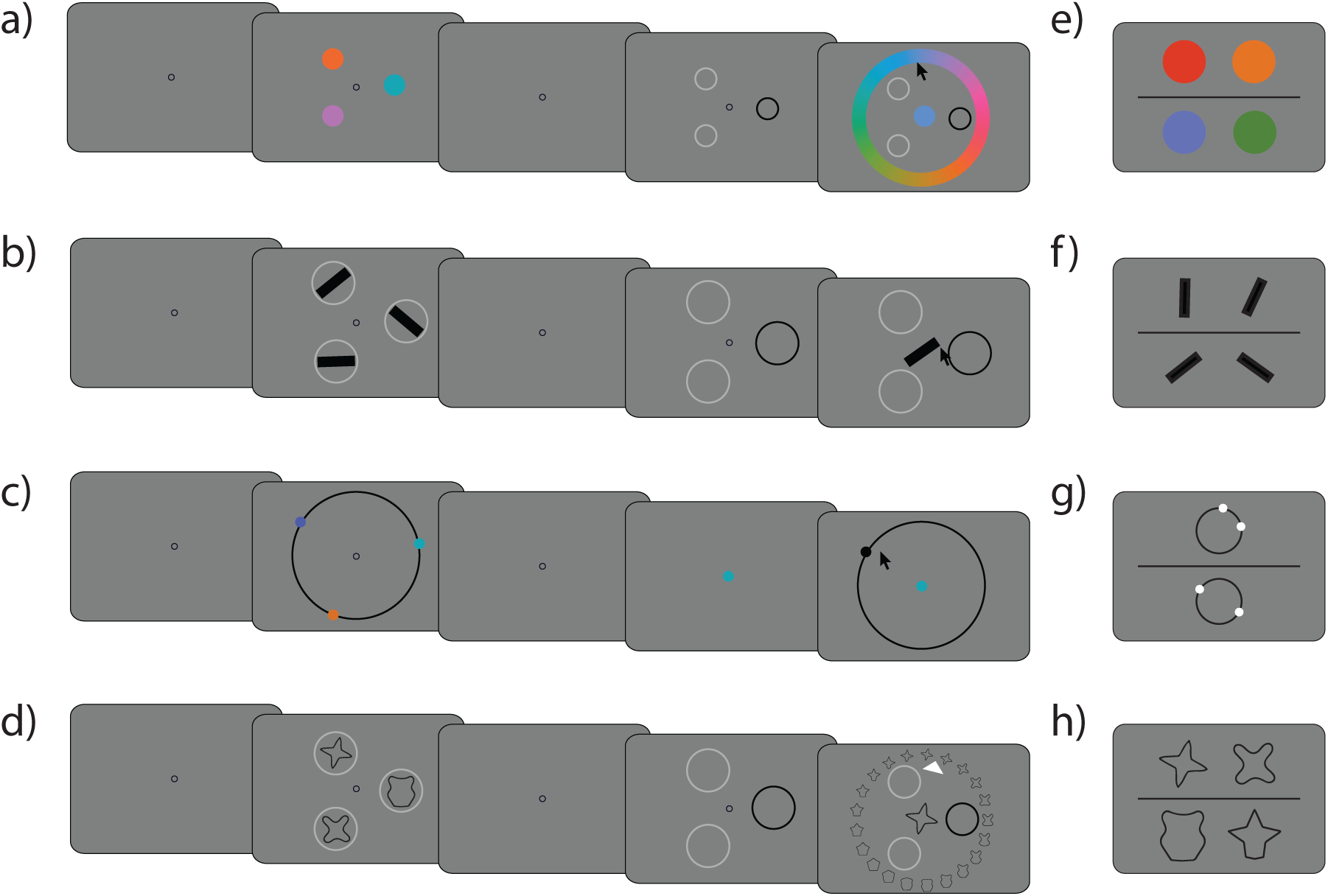
Procedure of experiments. *Note*. **(a-d)** Working memory analogue report task. On each trial, three or six objects were briefly presented and observers were required to memorize relevant feature. After a brief delay, one item was indicated with a recall cue and observers were asked to reproduce the memorized feature value on a continuous scale. **(e-h)** Psychophysical scaling quad task. Observers were asked to judge which of the two presented pairs (top or bottom) consisted of less similar stimuli. **(a & e)** color. **(b & f)** Orientation. **(c & g)** Angular location. **(d & h)** Shape. Stimuli not drawn to scale.

#### Color

The color stimuli were samples drawn from a color wheel, defined as a circle in CIELAB space with constant luminance (*L*\*= 50), centered at *a*\* = *b*\* = 20, with radius = 60 units and converted to RGB for presentation. In all tasks, color stimuli were presented as colored disks with a radius of 50 pixels.

#### Orientation

Orientation stimuli were randomly oriented black bars (100 × 15 pixels) selected from the unique orientation space [0°, 180°).

#### Angular location

In the angular location study, stimuli were disks positioned on a visible circle. In the working memory task, disks were colored by randomly sampling without replacement from nine salient colors (red, light blue, dark blue, cyan, green, dark green, purple, pink, orange). When more than one location stimulus was presented, locations were chosen at random with a minimum separation between disks of 5° of angular distance. In the perceptual analogue report task all disks were black. In the quad task, stimuli were white disks (3 pixels), presented in pairs on separate black rings (radius 100 pixels).

#### Shape

The shape stimuli were 360 unique shapes forming a circular space whose perceptual uniformity has been empirically validated (A. Y. Li, Liang, Lee, & Barense, 2020). Shape stimuli across all tasks were 130 × 130 pixels in size.

### Working memory analogue report task

Each trial started with the presentation of a central annulus (radius 5 pixels) (Figure 2 a-d). After 750 ms a memory sample array consisting of three or six stimuli was presented for 500 ms (1000 ms for shape due to greater stimulus complexity). This was followed by a delay period lasting 1000 ms and a probe display indicating one of the stimuli to be reproduced from memory.

In the color, orientation, and shape tasks (Figure 2 a, b, d), each stimulus was presented at one of six equidistant locations on an imaginary circle (radius 250 pixels). The probe display consisted of placeholder annuli at each of the previously occupied locations, all light-gray except for a single black placeholder indicating to observers that they should report the feature of the item previously at that location. Once a mouse movement was registered, a central stimulus appeared which observers could adjust by moving the cursor. For orientation stimuli, the central stimulus rotated with movements of the cursor. For color and shape stimuli, observers moved their cursor over a response wheel surrounding the probe display (Figure 2 a & d). Response wheels were rotated randomly from trial to trial. In the angular location task (Figure 2 c), the probe display consisted of a centrally presented disk in the same color as one of those previously shown, indicating which item’s location should be reported. Once a mouse movement was registered, a black ring and a black disk randomly positioned on that ring were displayed; the location of the disk could be adjusted using the mouse. For all stimulus features, observers registered their response with a mouse click. Each observer completed 15 practice trials followed by 100 trials which were submitted to analysis.

### Perceptual analogue report task

On each trial, a single stimulus was presented and remained visible while observers adjusted a central stimulus to match it. For the color, orientation, and shape reproduction task, the presentation and response phases were otherwise identical to the working memory task. In the location reproduction task, a single black disk (radius 3 pixels) was shown on every trial on a black annulus (radius 250 pixels) positioned at the centre of the screen. Once a mouse movement was registered, another randomly smaller (radius 210 pixels) or larger (radius 290 pixels) black annulus was centered on the same location along with a black disk whose location observers could adjust using the mouse. For all features, observers completed 10 practice trials followed by 50 trials which were submitted to analysis.

### Psychophysical scaling quad task

In order to establish the relationship between the physical and perceived distances in a stimulus dimension we used a psychophysical scaling “quad” task (Figure 2 e-h). On each trial, observers saw two pairs of stimuli, located above and below a central horizontal line, and were asked to judge which pair consisted of stimuli that were less similar.

For all stimulus features, the entire feature space was discretized in steps of 5° (2.5° for orientation), for a total of 72 unique stimuli. On every trial, four circular angles and associated feature values were sampled pseudorandomly from this discretized space. The first two stimuli, *θ*_1_ and *θ*_2_, were sampled randomly without replacement and with the constraint that the signed circular distance between them was smaller or equal to 180° (90° for orientation). The remaining two stimuli, *θ*_3_ and *θ*_4_, were sampled with the same constraint from the remainder of the circular space to avoid overlapping intervals. On each trial, coin-toss randomization was performed to determine the placement of the first pair (above or below the vertical centre of the screen).

In the color, orientation, and shape tasks (Figure 2 e, f, h) the four stimuli on each trial were centred on one of four corners of an imaginary rectangle positioned at the centre of the screen (side 160 pixels). In the angular location task (Figure 2 g) the stimuli were presented in pairs on two black rings centered in the upper and lower part of the screen (vertical offset 120 pixels). Observers responded by pressing the up or down arrow-key to report that stimuli in the top or bottom pair were less similar, respectively. The trials were not speeded, and the stimuli remained visible until observers chose an option. Observers completed 210 trials for colour, 230 trials for orientation, and 240 trials for location and shape features. The number of trials was chosen based on pilot data to produce an average testing block duration of approximately 15 minutes. Before analysing the data, we excluded all trials on which the response time was faster than 200 ms (less than 0.1% of trials for all features).

## Analysis

For ease of comparison, all stimulus values were analysed and are reported with respect to the circular parameter space of possible values, [−*π, π*) radians. To quantify the degree of evidence supporting an effect, we used Bayesian hypothesis tests, implemented in JASP (JASP Team, 2020) with the default Jeffreys-Zellner-Siow prior on effect sizes (Liang, Paulo, Molina, Clyde, & Berger, 2008). The Bayes factor compares the relative predictive adequacy of two competing hypotheses (e.g., alternative and null) and quantifies the change in belief that the data bring about for the hypotheses under consideration (Wagenmakers et al., 2018). Values of BF_10_ *>* 1 indicate evidence for an effect and values BF_10_ *<* 1 indicate evidence for the absence of an effect. Results of Bayesian ANOVA are reported as overall evidence for an effect, BF_*incl*_, which quantifies how much more likely is the data under models that include the effect of interest compared to models that do not include the same effect (Hinne, Gronau, van den Bergh, & Wagenmakers, 2020). We interpret the values of Bayes factors according to a classification scheme as in Lee and Wagenmakers (2013): BF = 1 as no evidence; 1 *<* BF *<* 3 as anecdotal evidence; 3 ≤ BF *<* 10 as moderate evidence; 10 ≤ BF *<* 30 as strong evidence; 30 ≤ BF *<* 100 as very strong evidence; BF ≥ 100 as extreme evidence. Unless specified otherwise, correlations were calculated after removing outlying data points iteratively until all data fell within ±3 SD of the mean (average of 1.7% of excluded data per measure, range 0.5% - 3.8%). When a correlation is reported for data pooled across different features, the data for each stimulus feature was first standardized and outliers removed before pooling the data and calculating the effect of interest.

### Psychophysical similarity function

We applied a similarity-scaling technique to data collected on the quad task. We did this by adapting the Maximum Likelihood Difference Scaling (MLDS) method proposed by Maloney and Yang (2003). Given a set of physical distances {*ψ*_1_, *ψ*_2_, …, *ψ*_*k*_} in the feature space, the goal of the MLDS method is to find a set of psychophysical distance values {Ψ_1_, Ψ_2_, …, Ψ_*k*_} that best predict an observer’s judgments about stimulus similarity.

On every trial we presented two pairs of stimuli with physical feature values of *θ*_*a*_, *θ*_*b*_, *θ*_*c*_ and *θ*_*d*_, giving respective physical distances *ψ*_*ab*_ = |*θ*_*a*_ ⊖ *θ*_*b*_| and *ψ*_*cd*_ = |*θ*_*c*_ ⊖ *θ*_*d*_|, where ⊖ is a subtraction on a circle. We asked observers to judge which pair consisted of stimuli that were less similar to each other. If we denote the perceived difference within each pair as Ψ_*ab*_ and Ψ_*cd*_, and assuming the difference estimates are corrupted by normally distributed additive noise, observers should judge the pair *ab* as having the larger difference when Ψ_*ab*_ *>* Ψ_*cd*_ + *ϵ*, where *ϵ* ∼ *N* (0, *σ*^2^).

The only constraint in the above description is that {Ψ_1_, Ψ_2_, …, Ψ_*k*_} change monotonically, however empirical studies have found that perceived similarity is approximately exponential with distance in feature space across a wide range of sensory modalities and stimulus features (Schurgin et al., 2020; Sims, 2018). Here, we leveraged that knowledge and constrained {Ψ_1_, Ψ_2_, …, Ψ_*k*_} to decay exponentially with distance. The similarity function *ω* replaces a set of psychophysical distances {Ψ_1_, Ψ_2_, …, Ψ_*k*_} and is defined as:

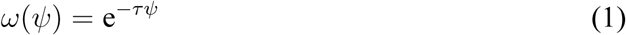

This parametric description reduces the number of free parameters to two: *τ* which corresponds to the decay rate of the distance function and the SD parameter *σ* which reflects the stochasticity in the decision model.

Empirical similarity functions display a small but systematic deviation from exponential which is thought to be due to variability in perception of the individual stimuli, rather than distances between stimuli (Nosofsky, 1992; Schurgin et al., 2020). To account for this, we convolved the exponential similarity function with perceptual noise estimated from the perceptual analogue report task:

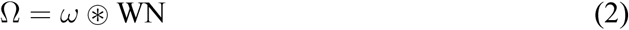

where Ω is the final form of the similarity function, and WN is a wrapped normal distribution with mean of zero and standard deviation matched to errors observed in the perceptual analogue report task. Schurgin et al. (2020) found this parametric description provided an excellent fit to similarity judgment data. Finally, the similarity function was normalized to the range [0, 1]. The similarity function Ω was fit to each observer’s data and the best fitting parameters *τ* and *σ* were found using maximum likelihood estimation.

### Perceptual analogue report

Errors in the perceptual analogue report task were used to derive estimates of perceptual noise. Data was pooled across observers and errors that were *>*3 SD away from the mean were excluded. The SD of the best fitting zero-mean wrapped normal distribution was found via maximum likelihood search.

### Working memory

We fit three models to recall errors from the WM analogue report task: the Neural Resource model (Fig. 1a-d), the empirical TCC model using the similarity function estimated via the parametric MLDS method (Fig. 1f-h), and a synthetic TCC model in which the similarity function was fitted to the WM data.

#### Neural Resource model

We fit the Neural Resource model of Bays (2014) to each observer’s responses on the working memory task (Fig. 1a-d). In this model of WM based on population coding, memory stimuli are encoded in the noisy firing activity of feature-selective neurons, and recall estimates are obtained by optimal decoding of the population activity over a fixed time window. The idealized neurons have homogeneous von Mises tuning functions that evenly tile a circular feature space. In a population of *M* neurons, the average response of the *i*th neuron with preferred value *φ*_*i*_ in response to a stimulus value *θ* is given by

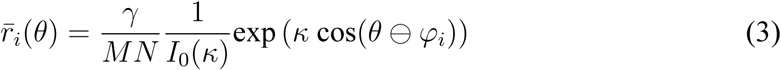

where *γ* is the total population activity, *N* is the number of memorized items, *κ* is the von Mises tuning width, and *I*_0_(·) is the modified Bessel function of the first kind with order zero. Equation 3 implements a simple form of divisive normalization (Carandini & Heeger, 2011) whereby the total spiking activity is divided between the items in memory. The probability of the *i*th neuron producing *n*_*i*_ spikes in the interval *T* is modeled as a homogeneous Poisson process:

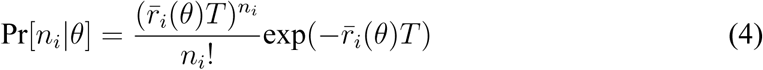

The reported value for the cued item is obtained by maximum likelihood decoding of the population spiking activity, **n**:

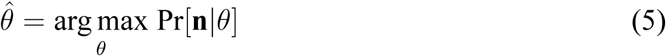

The parameterization in terms of *γ* and *κ* fully specifies the population response to a stimulus and hence the distribution of recall errors predicted under the model. However, with this parameterization, the responsiveness function’s height and width are not independently specified, as the *κ* parameter affects both features of the function. To facilitate comparison with the TCC model, we reparameterized the model (as previously described by Bays & Taylor, 2018) in terms of the peak firing rate (*r*_*max*_) and the full-width at half-maximum of the tuning function (*W*). Note that this reparameterization does not affect the predictions of the model or the quality of fits.

We calculate *r*_*max*_ by noting that the neuron’s maximum firing rate is achieved for a stimulus that matches the neuron’s preferred value, i.e., when *θ* = *ϕ*_*i*_, which leads to:

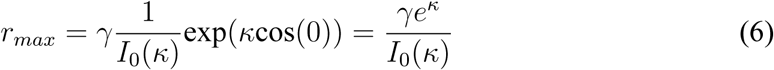

Similarly, half-maximum points lie symmetrically on either side of *ϕ*_*i*_, and *W* corresponds to the distance between these points:

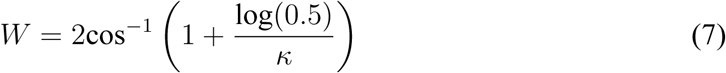

The peak activity parameter *r*_*max*_ determines each neuron’s response to a visual stimulus matching its preferred value. As *r*_*max*_ increases, more spikes are available for the decoding of feature value, resulting in smaller deviations between the true and reconstructed feature value. The *W* parameter controls the range over which feature values deviating from the preferred value elicit a response. This primarily affects the shape of the resulting error distributions.

Maximum likelihood fits were obtained via the Nelder-Mead simplex method (function *fminsearch* in Matlab). A MATLAB toolbox implementing the neural population model is available to download from https://bayslab.com/toolbox.

#### The Target Confusability Competition model

##### Empirical-TCC

In the TCC model of WM proposed by Schurgin et al. (2020), the distribution of recall errors depends on a perceptual similarity function Ω, which is a non-linearly decreasing function of distance in the stimulus space (Fig. 1e-h). On recall of a stimulus with true feature value *θ*, a noisy memory-match signal is generated for each of a set of possible feature values *θ*^*′*^, as a draw from a unit-variance Gaussian distribution:

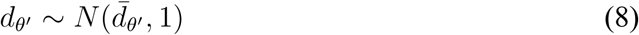

The mean 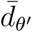 of the memory-match signal distribution is determined by the psychological similarity of each feature value *θ*^*′*^ to the true stimulus value *θ*, through a function Ω:

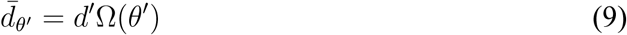

where *d*^*′*^ is a free parameter. The reported feature value is the one with the strongest memory-match signal (a “winner-takes-all” decision rule):

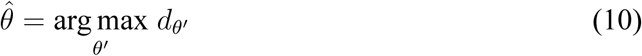

We followed Schurgin et al. (2020) by assessing *d*_*θ*_*′* at 360 equally spaced points in the stimulus space, and adding a fixed normally distributed error to recall estimates (SD = 2°) intended to represent motor error. We employed a grid search to obtain maximum likelihood model fits. The *d*^*′*^ parameter was evaluated on a dense grid ranging from 0 to 5.5 in increments of 0.02.

##### Synthetic-TCC

In the original version of TCC, the responsiveness function Ω is estimated from a separate psychophysical distance task such as a quad task, under the assumption that the pattern of WM errors derives from the psychological similarity between features. To test this assumption, we compared responsiveness functions obtained in this way to synthetic responsiveness functions chosen to best fit the working memory data. Mirroring the parametric MLDS method described in the main paper, we modelled the synthetic responsiveness function as an exponential convolved with perceptual noise estimated from the perceptual analogue report task, with the exponent as a free parameter. Along with the *d*^*′*^ parameter, this made the extended TCC model a two-parameter model. When fitting across set sizes, we fit separate *d*^*′*^ parameters but searched for a single best fitting exponent across both set sizes, making three free parameters in total. To find the best-fitting synthetic function, we used a bank of exponential functions with the exponents ranging from 0.001 to 10 in steps of 0.0313.

##### von Mises-TCC

In addition, we fit a version of the Synthetic-TCC model with a von Mises responsiveness function instead of the exponential-like function. We again searched for the best fitting responsiveness function across two set sizes, on a grid of circular standard deviation from 0.03 to 2 in steps of 0.0062.

#### Relating the models

Here we briefly clarify the mathematical correspondence between TCC and Neural Resource models (previously presented in a preliminary form in Bays (2019)). We start by simplifying the equation for mean firing rate in the Neural Resource model (Eq. 3):

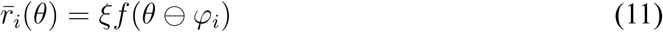

Where 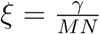 is the response amplitude and *f* (·) is the von Mises function. We then rewrite the TCC Equation 9 as:

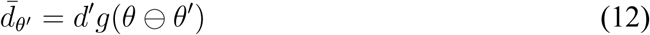

where *g*(·) is a non-linearly decreasing similarity function. It becomes clear that the mean spiking response 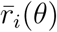 in Eq. 11 corresponds to the mean memory-match signal 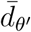 in Eq. 12, with *ξ* and *d*^*′*^ scaling the activation of the respective responsiveness function.

In TCC, the response to a stimulus *θ* is drawn from a unit-variance Gaussian distribution (Eq. 8). Using a Gaussian approximation to Poisson for the Neural Resource model (as in e.g. Schneegans and Bays 2017), neural responses can be modelled as:

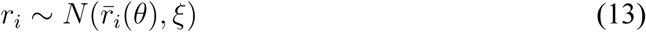

In the Gaussian case the neural signal can be scaled arbitrarily without changing predictions of the model as long as the ratio of signal to noise remains unchanged. Scaling the response function by a factor 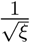 results in a unit-variance Gaussian 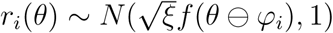. Finally, setting 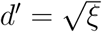 makes the distributions of signal strengths in the two models identical. Critically, this correspondence reveals that the main difference between the two models lies in the choice of the responsiveness functions *f* (·) and *g*(·), which is the topic of the main paper.

The TCC model does not specify how *d*^*′*^ varies with set size, however the relationship identified here between the signal detection variable *d*^*′*^ and response amplitude *ξ* in the Neural Resource model leads to the prediction that *d*^*′*^ is proportional to 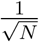, which we note is also consistent with predictions of the sample-size model of Shaw (1980), indicative of the deep theoretical connections between these models (Schneegans et al., 2020; Smith, 2015).

#### Correction for attenuation

To assess the correspondence between the empirical similarity functions used by TCC and the best-fitting exponential activation functions found in the synthetic-TCC model, we calculated the Pearson correlation coefficient between the exponents of two functions. As the magnitude of this correlation can be reduced due to measurement error of one or both measures, we applied a correction for attenuation using Spearman’s 1904 formula:

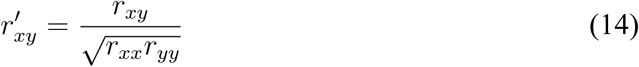

where *r*_*xy*_ is the observed correlation calculated from the raw data, and *r*_*xx*_ and *r*_*yy*_ are the reliabilities of measurements *x* and *y*, respectively. Estimates obtained by applying this correction are necessarily larger than the uncorrected correlation.

We estimated the reliabilities of both measures using a bootstrap method. In particular, for each observer we generated pairs of empirical exponents and pairs of synthetic exponents by resampling with replacement the trial-by-trial data from the quad and WM tasks, respectively. We repeated this procedure to generate 100 of these simulated samples per observer. As in our main correlational analysis, observations that were ±3SD away from the sample’s mean were excluded. The reliability was estimated as the mean correlation across all simulated samples. This method was repeated for each feature separately.

## Results

We measured participants’ ability to reproduce visual stimuli from perception and working memory in four different feature dimensions using standard analogue report designs. In the same participants, we assessed psychological similarity functions for each feature dimension using the method of quadruples, in which observers were required to compare two pairs of stimuli and decide which pair consisted of less similar stimuli.

### Psychophysical similarity functions

We employed a parametric version of the MLDS method (Maloney & Yang, 2003) to fit empirical similarity judgments, obtaining two maximum likelihood parameters for each participant: an exponent *τ* that describes how quickly similarity decays with distance in the feature space, and a standard deviation *σ* which reflects variability in the distance comparisons but does not contribute to the distance function itself. To account for effects of perceptual noise on individual stimuli, the exponential function was convolved with a wrapped normal distribution matched in SD to the corresponding perceptual reproduction task (0.103, 0.147, 0.022, 0.137 for color, orientation, location, and shape, respectively) and then scaled to the range [0, 1]. The estimated similarity functions, shown in Fig. 3a, are consistent in shape with functions previously reported using non-parametric methods (Nosofsky, 1992; Schurgin et al., 2020).

**Figure 3.**
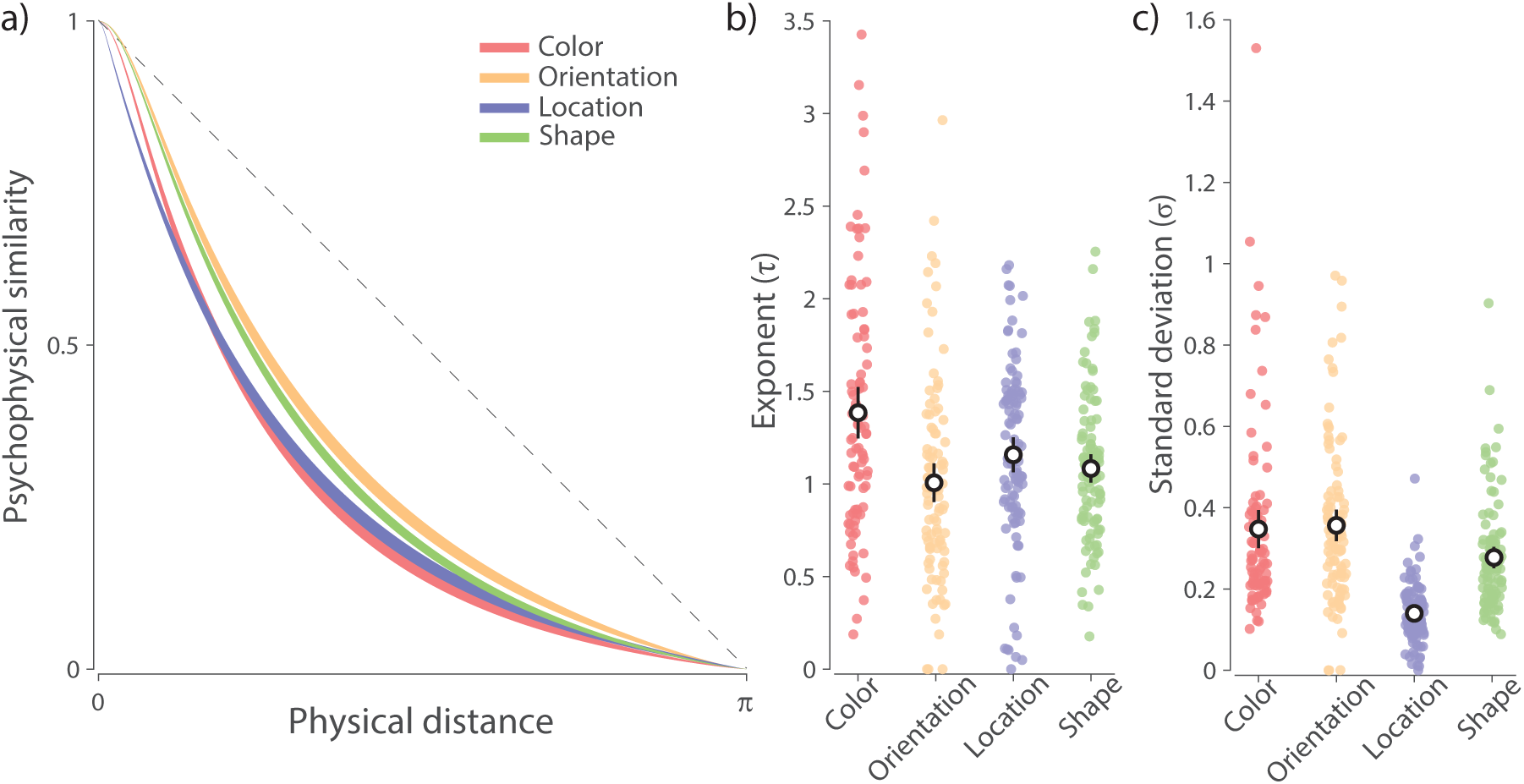
Perceptual similarity estimates.

*Note*. **(a)** Similarity functions estimated via the parametric MLDS method (mean ± SE) from judgments in the quad task. The dashed black line corresponds to a linear relationship between similarity and physical distance in the feature space. Larger divergence from dashed line indicates greater non-linearity in the perceptual space. **(b-c)** Best fitting MLDS parameters. Colored dots show individual observers’ fitted parameter values and the black circles with errorbars show the mean and 95% credible intervals. **(b)** Exponent parameter. Higher values correspond to more rapid decay of similarity with physical distance. **(c)** Standard deviation parameter. Higher values indicate greater variability in similarity judgments.

Bayesian ANOVA applied to the fitted exponent parameters (Fig. 3b) found extreme evidence for a difference in the curvature of the similarity functions across features (BF_*Inclusion*_ = 1415). Comparing each pair of features, we found color displayed the largest non-linearity (all BF_10_ ≥ 4.7). The remaining three features displayed similar non-linearities, as indicated by weak evidence for a difference in curvature (location vs. orientation: BF_10_ = 1.2), or moderate evidence against a difference in curvature (shape vs. location and orientation: all BF_10_ ≤ 0.29). For the noise parameter (Fig. 3c), there was extreme evidence for differences between feature dimensions (BF_*Inclusion*_ = ∞), with location displaying the lowest noise estimates (all BF_10_ ≥ 7.4 × 10^12^). Shape was found to have lower noise estimates than color and orientation (all BF_10_ ≥ 3.8), while color and orientation displayed comparable levels of noise (BF_10_ = 0.16).

### Working memory performance

Data points in Figure 4a show error distributions from the working memory analogue report task for each of the four feature dimensions. As typically observed, increasing the number of memory items increased recall variability for all features, seen here as a greater dispersion of errors for set size 6 (gray dots) compared to set size 3 (black dots). We also observed qualitative differences in recall variability between the different feature dimensions. In line with previous studies, the Neural Resource model (Fig. 1) provided a very good description of empirical error distributions in each feature dimension, capturing the effects of set size and the leptokurtosis (“long tails”) of error distributions. The mean predictions of the model are shown as colored lines in Figure 4a, and best-fitting model parameters are plotted in Figure 4c.

**Figure 4.**
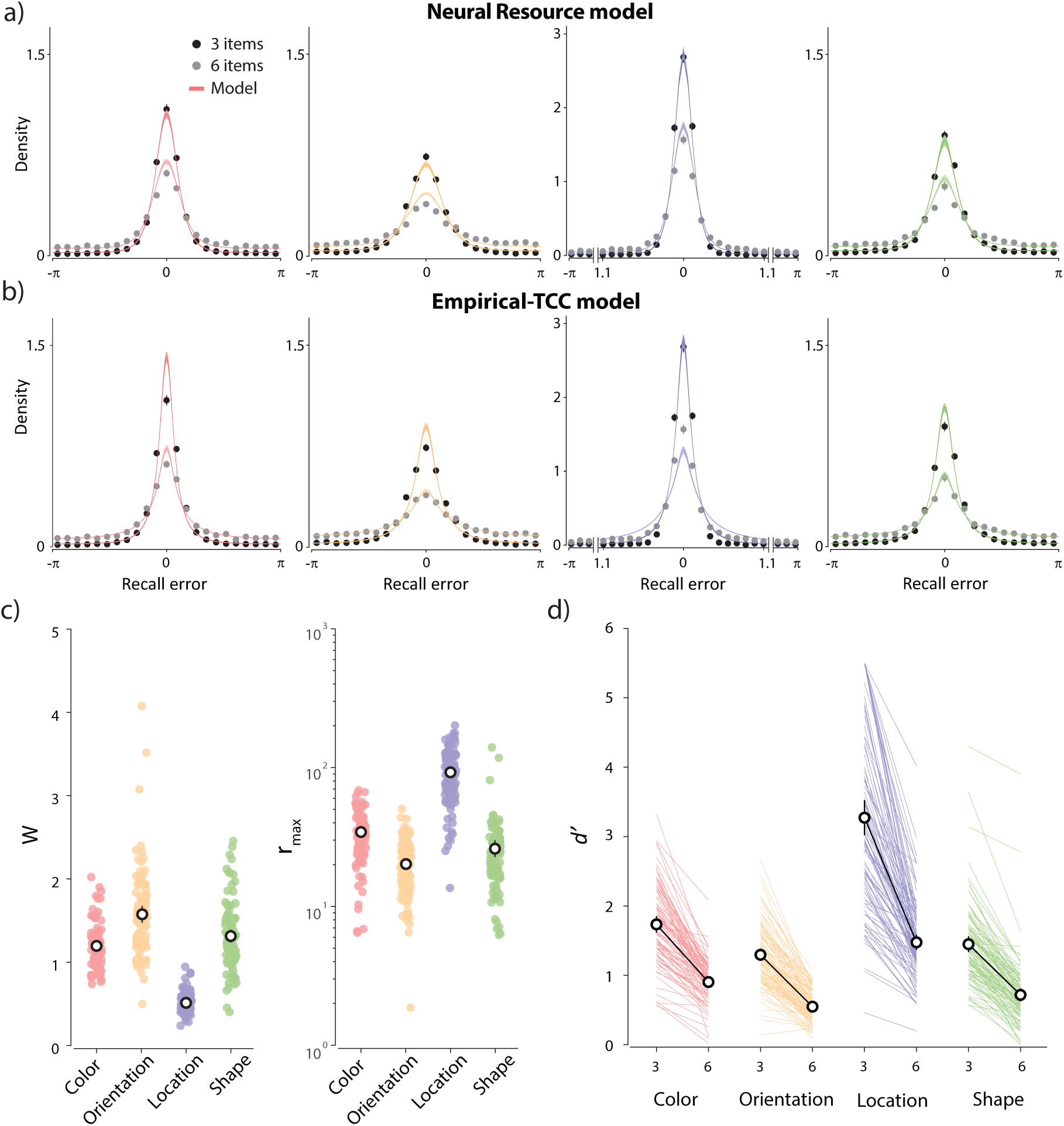
WM error distributions and fits of the Neural Resource model and the TCC model based on empirical similarity functions. *Note*. **(a-b)** Black and grey data points and errorbars show recall errors (mean ± SE) for set sizes three and six, respectively. Colored curves and patches show predictions of the models (mean ± SE). Note that, for angular location (blue), the central region of the x-axis has been rescaled for better visibility. **(a)** The Neural Resource model fit. **(b)** The TCC model fit using empirical similarity functions. **(c)** The best fitting tuning width (*W*) and peak firing rate (*r*_*max*_) parameters from the Neural Resource model. **(d)** The best fitting memory-match signal (d^*′*^) parameters from the TCC model for each set size. **(c-d)** The black circles with errorbars show the mean and 95% credible intervals.

Colored lines in Fig. 4b show fits to the same data of the TCC model based on empirical similarity functions, derived by applying the parametric MLDS method to the same participants’ responses in the quad task (Fig. 3a). For color, orientation, and shape datasets, the TCC model qualitatively captured the error distributions, including the decline in recall precision with set size. The latter was achieved by fitting separate *d*^*′*^ parameters to each set size (Fig. 4d), since the TCC model, unlike the Neural Resource model, does not prescribe how the memory-match signal changes with number of memory items. For the angular location dataset (blue in Fig. 4b), TCC qualitatively failed to reproduce the observed recall errors. Comparing the TCC and Neural Resource fits with the AIC metric, we found extremely strong evidence in favour of the population model for location (mean ΔAIC = 15.57, t-test on individual differences in AIC values: BF_10_ = 7.65 × 10^8^), weak evidence in favour of the population model for color (mean ΔAIC = 1.57, BF_10_ = 1.55) and moderate evidence against an advantage for either model for orientation (mean ΔAIC = 0.49, BF_10_ = 0.17) and shape (mean ΔAIC = -0.08, BF_10_ = 0.11).

### Comparing empirical and synthetic similarity functions

The differences in quality of fit between the Neural Resource and TCC models could in principle arise from differences in the detailed architecture of the models (Fig. 1), or due to the TCC model employing empirical psychophysical similarity as the basis of its responsiveness function. To distinguish these possibilities, we evaluated a synthetic version of the TCC model in which we fit the curvature of the similarity function to the WM data along with the memory-match signal parameters. Unlike the Empirical-TCC model, this model provided a satisfactory fit to all four feature dimensions. Despite being penalized for its greater complexity, the Synthetic-TCC model showed a substantial improvement in AIC compared to Empirical-TCC for location (mean ΔAIC = 14.6, BF_10_ = 4.14 × 10^9^) and, to a lesser degree, for color (mean ΔAIC = 1.79, BF_10_ = 15.66) and orientation (mean ΔAIC = 1.26, BF_10_ = 16.88). Finally, the fits for the shape dataset were comparable across the two versions of TCC (mean ΔAIC = 1.14, BF_10_ = 0.57).

These results indicate that TCC has an adequate architecture to fit WM data; however, basing the tuning function on psychophysical similarity estimated for the same feature space appears suboptimal. Comparing empirical similarity functions estimated from the quad task (Fig. 3a) and the synthetic similarity functions that best fit WM data (Fig. 5b), we found very strong evidence for a difference for location (BF_10_ = 2.51 × 10^20^), with the synthetic exponent parameters being larger on average (Fig. 5c). The opposite pattern was observed for both color and orientation, with strong evidence that empirical exponents overestimated the steepness of the WM response function (color: BF_10_ = 4001; orientation: BF_10_ = 37.12). Finally, we found moderate evidence against differences between empirical and synthetic similarity curves for the shape feature dimension (BF_10_ = 0.11).

**Figure 5.**
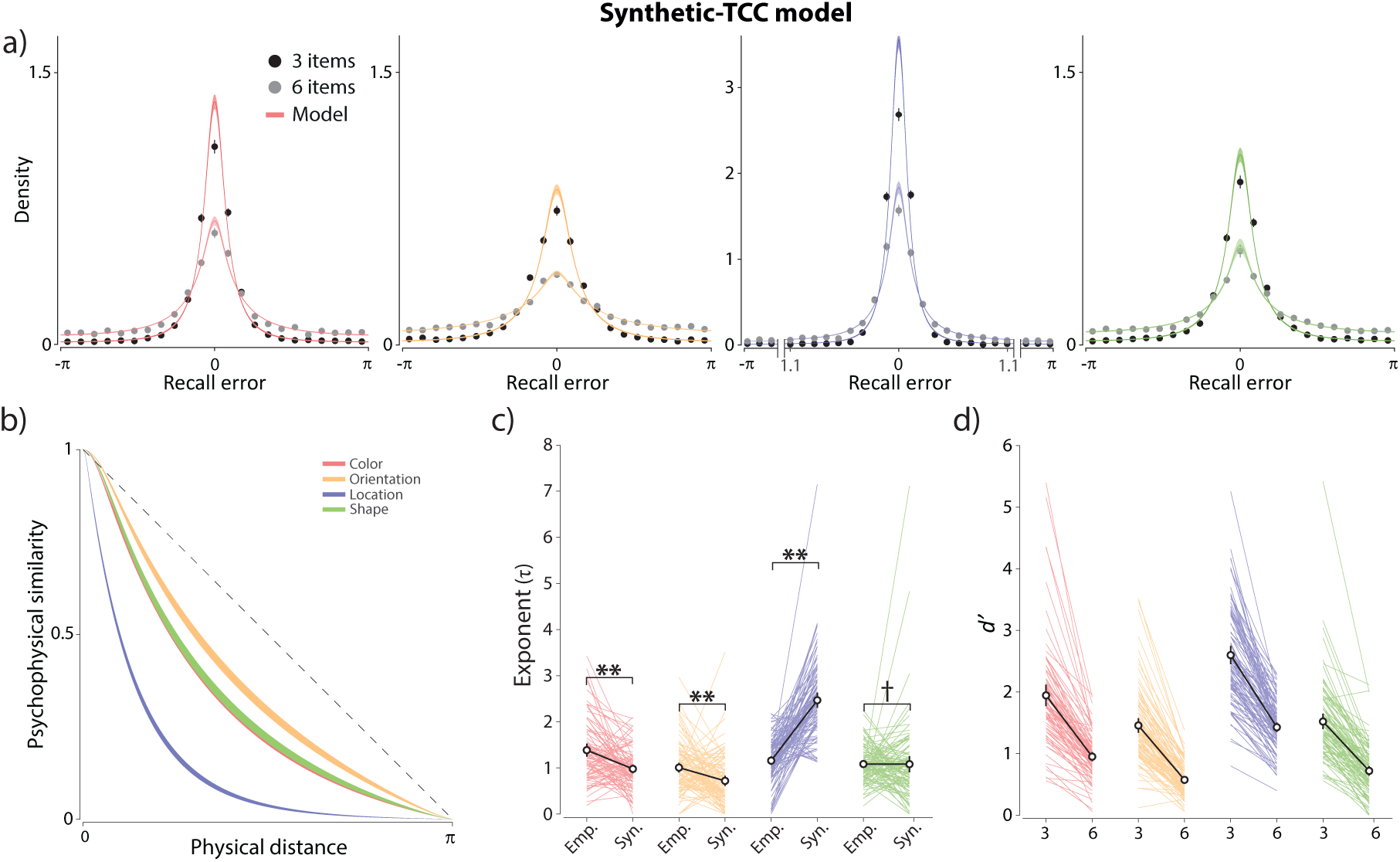
Fits of the TCC model with synthetic similarity function to WM errors. *Note*. **(a)** The Synthetic-TCC model fit. **(b)** The best fitting synthetic similarity functions derived from WM data. **(c)** Comparison of empirical exponent parameters obtained from the perceptual similarity task and synthetic exponent parameters that best fit WM data. **(d)** The best-fitting memory-match signal (d^*′*^) parameters from the Synthetic-TCC model. Double asterisks indicate strong evidence for a difference (BF_10_ *>* 10) and single dagger indicates moderate evidence against a difference (BF_10_ *<* 0.3). **(c-d)** The black circles with errorbars show the mean and 95% credible intervals.

Next, we looked for correlations between parameters of the MLDS fit describing a participant’s responses on the quad task, and parameters of the Neural Resource and Synthetic-TCC models fit to the same participants’ WM data, with a specific focus on the parameters that contribute to the shape of the WM error distributions. If the psychophysical similarity function was the basis for the shape of WM error distribution, we would expect to see a correlation between the curvature of the empirical similarity function derived by the MLDS method and the best fitting responsiveness functions estimated using WM data. In contrast to this prediction, we observed moderate evidence for absence of a correlation between the exponents of the empirical similarity functions and the synthetic similarity functions that reproduced WM errors (r(387) = −0.07, BF_10_ = 0.16; Fig. 6a). Indeed, we found moderate evidence for absence of a correlation for each of the four feature dimensions: color r(89) = 0.07, BF_10_ = 0.16; orientation r(96) = −0.13, BF_10_ = 0.28; angular location r(103) = −0.14, BF_10_ = 0.32; shape r(99) = −0.06, BF_10_ = 0.15. We confirmed that the absence of correlations could not be explained by measurement noise in the estimated exponents. Reliability estimates for the empirical exponents were 0.61, 0.59, 0.54, and 0.54 for color, orientation, location, and shape, respectively. For the synthetic exponents we found comparable reliabilities of 0.57, 0.53, 0.68, and 0.51 for color, orientation, location, and shape, respectively. Correcting the correlations for attenuation based on these reliabilities (see Methods) still yielded no meaningful associations between the empirical and best-fitting exponents: color r = 0.12; orientation r = −0.23; angular location r = −0.23; shape r = −0.11. Across all four features, the uncorrected association accounted for at most 2.6% of variance and the corrected association for at most 5.4%.

**Figure 6.**
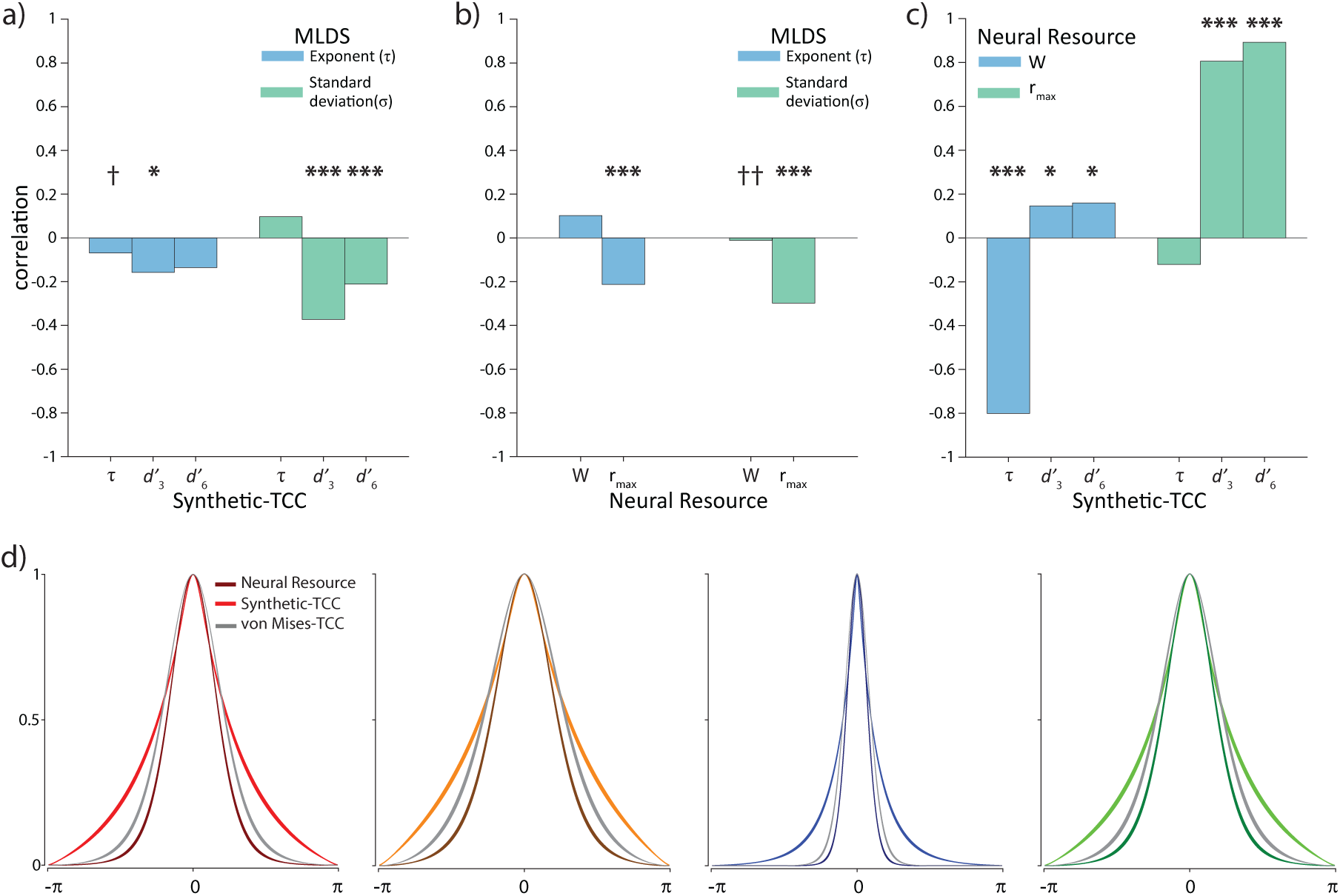
Correlation between models’ parameters, and the best-fitting activation functions. *Note*. **(a-c)** Correlations between the parametric MLDS, Synthetic-TCC, and Neural Resource model parameters. Asterisks indicate evidence for a correlation (* BF_10_ *>* 3; *** BF_10_ *>* 100), and daggers indicate evidence against a correlation († BF_10_ *<* 0.3; †† BF_10_ *<* 0.1). **(d)** The best-fitting activation functions (mean ± SE) from the Neural Resource, Synthetic-TCC, and von Mises-TCC models for (L to R) color, orientation, angular location, and shape.

Similarly, the curvature of the empirical similarity function did not correlate with the tuning width parameter that determines the shape of error distributions in the Neural Resource model (r(387) = 0.1, BF_10_ = 0.47) (Fig. 6b). Together, these results provide strong evidence that the empirical similarity function estimated from the perceptual quad task is unrelated to the shape of the WM error distribution.

In contrast, the MLDS exponent parameter was related to the remaining parameters of Synthetic-TCC 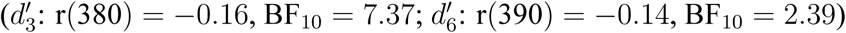 and Neural Resource models (r_*max*_: r(390) = −0.21, BF_10_ = 494). However, once the effect of the MLDS noise parameter was partialled out, we found weak to strong evidence for absence of these correlations (all BF_10_ ≤ 0.43). Moreover, we did not observe a meaningful association between the MLDS noise parameter and the exponent from the Synthetic-TCC model (r(375) = 0.09, BF_10_ = 0.373) or the width parameter from the Neural Resource model (r(374) = −0.01, BF_10_ = 0.07).

We found extremely strong evidence for a correlation between the MLDS noise parameter and the signal amplitude parameters of the model, both for Synthetic-TCC 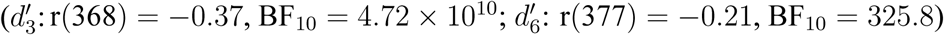 and Neural Resource (r_*max*_: r(377) = −0.3, BF_10_ = 2.13 × 10^6^). The negative sign of these correlations indicates that observers who provided noisier judgments of perceptual similarity in the quad task also reported memoranda less precisely in the WM analogue report task. This association indicates a shared source of variability across tasks, potentially related to broader cognitive ability or stable levels of engagement in cognitive tasks.

### Relationship between Neural Resource and TCC parameters

The exponent in the Synthetic-TCC model, which was fitted to WM data, was strongly correlated with the tuning width parameter in the Neural Resource model (r(382) = −0.80, BF_10_ = 1.46 × 10^83^) (Fig. 6c). Similarly, the signal-strength parameters d’ for each set size in the Synthetic-TCC model were strongly correlated with the signal amplitude parameter r_*max*_ in the Neural Resource model (r_*max*_ and 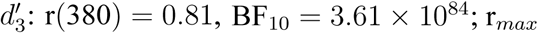 and 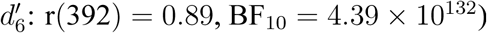). These results are consistent with the close correspondence in architecture between the two models.

Finally, the correlation of the signal amplitude parameters from the Synthetic-TCC model and the tuning width parameter in the Neural Resource model received moderate support 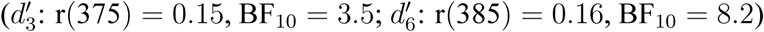, while no evidence was found for correlation of the peak firing rate and the exponent from the Synthetic-TCC model (r(387) = −0.12, BF_10_ = 1.1).

Double asterisks indicate strong evidence for a difference (BF_10_ *>* 10) and single dagger indicates moderate evidence against a difference (BF_10_ *<* 0.3).

### Synthetic-TCC with von Mises responsiveness function

Comparing the responsiveness functions of the fitted Synthetic-TCC and Neural Resource models (Fig. 6d; light and dark colored lines, respectively) we observed a strong qualitative similarity in the vicinity of the functions’ peak, suggesting that the particular combination of exponential scaling convolved with perceptual noise specified by TCC partially emulated the von Mises (circular normal) tuning function assumed by the Neural Resource model. Having established this correspondence, we next tested whether the exponential-like activation function used by TCC provided a better account of WM data than the von Mises function typically used to approximate neural tuning functions. To this end we fit a version of the Synthetic-TCC model with a von Mises responsiveness function. This model showed a substantial improvement in AIC over the model with exponential-like responsiveness function for location (mean ΔAIC = 2.92, BF_10_ = 2.69 × 10^6^) and orientation (mean ΔAIC = 1.03, BF_10_ = 2325), while the fits were comparable for color (mean ΔAIC = 0.52, BF_10_ = 0.55) and shape (mean ΔAIC = 0.05, BF_10_ = 0.11). Responsiveness functions obtained from this model (Fig. 6d, gray lines) were notably similar to those obtained from the Neural Resource model.

## Discussion

We measured psychophysical similarity and working memory recall in four different stimulus spaces, to evaluate the hypothesized relationship between the psychological similarity function and the distribution of WM errors. Our findings provide clear evidence against such a relationship. At the group level, the responsiveness function that provided the best fit to working memory errors deviated strongly from the one obtained from psychophysical similarity measurements for three out of four stimulus dimensions. At the participant level, we observed evidence against a correlation between best-fitting and observed similarity functions in all four stimulus dimensions.

Having failed to corroborate a specific match between measured similarity functions and patterns of WM error in the same stimulus spaces, we further examined whether an exponentially decaying responsiveness function had a general advantage over the von Mises (circular normal) function more typical of population coding models. A version of TCC equipped with a von Mises responsiveness function fit the WM data equally well or better than the model based on an exponential function. The fitted von Mises responsiveness functions in this model were notably similar to the tuning functions obtained by fitting the Neural Resource model, an established model of WM based on population coding principles (Bays, 2014; Schneegans et al., 2020). This is consistent with the strong similarities in architecture between the models (Fig. 1).Furthermore, our findings on the advantage of the von Mises over the exponential responsiveness function are consistent with recent results reported by Oberauer (2021). By combining features of different descriptive models of WM within a space of possible models and fitting them to analogue report WM data, Oberauer demonstrated that a version of a model with a von Mises responsiveness function provides a consistently better fit compared to an equivalent model with the exponential responsiveness function.

Previous simulations of population coding have shown that the shape of the tuning function has a relatively small effect on patterns of decoding error (Bays, 2014; Bethge, Rotermund, & Pawelzik, 2002). Similarly, the long-tailed error distributions generated by the Empirical-TCC model are not primarily a result of a non-linear similarity function, but rather arise from the combination of noise distribution and decoding strategy, as in other population coding accounts. Indeed, long tails would be a feature of Empirical-TCC predictions even for a psychophysical similarity function that is veridical, i.e. linear with physical distance (Schurgin et al., 2020).

The source study showed that the TCC model with empirical similarity function fitted analogue report data better than some of the descriptive models widely used in the WM field (specifically, the normal-plus-uniform mixture model of Zhang and Luck (2008) and a variant of the variable precision model by Fougnie et al. (2012)). The study also showed that the TCC model could reproduce behavioural results from 2-AFC working memory tasks, where observers must distinguish a repeated stimulus from a foil. However, the study did not distinguish whether the relative success of TCC over the descriptive models was a consequence of incorporating an empirical similarity function into the WM model, or of the differences in model architecture that made it more similar to previous population coding accounts. In particular, the study did not systematically test whether variations in the empirical similarity function, across individuals or feature dimensions, were matched by corresponding changes in the shape of WM error distributions, nor were any alternative responsiveness functions compared to those obtained from the similarity judgment tasks.

Although we did not find the non-linearity of the similarity function to be related to WM errors, we found compelling evidence that the noise affecting judgments in the perceptual quad task correlated with the amplitude of the WM signal in both the Synthetic-TCC and Neural Resource models. Observers who provided less precise reports in the working memory task also discriminated pairs of stimuli in the quad task less accurately. A number of possible explanations for this effect should be considered. The common source of noise could indicate that the perceptual task in fact called upon working memory to facilitate the comparisons between stimuli(Woodman & Luck, 2004).

Alternatively, it could reflect an influence on both tasks of stable levels of engagement, or arise from differences between participants in broader cognitive ability (Ackerman, Beier, & Boyle, 2005). These possibilities might be disambiguated in future studies using purpose-designed experiments.

Leaving aside our main conclusion, there are many points of agreement between the claims made for TCC and those based on population coding and variable precision accounts (Fougnie et al., 2012; Schneegans et al., 2020; van den Berg et al., 2012). These include the demonstration that a purely continuous model is compatible with the long-tailed error distributions that have sometimes been taken as evidence for discrete representations (Adam, Vogel, & Awh, 2017; Zhang & Luck, 2008), and the important observation that a model with just one parameter is sufficient to capture variations in recall error distribution within a particular feature dimension (that parameter being *d*^*′*^ in the synthetic TCC model, activity amplitude *r*_*max*_ in Neural Resource, and mean precision 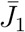 in the variable precision model). Comparison of the models also leads to a simple relationship between population activity amplitude and the signal detection parameter *d*^*′*^.

Besides difference in response function between the TCC and Neural Resource models, there are two other differences that relate to the implementation of the population code (Fig. 1). The first concerns the noise distribution, with TCC using a Gaussian and Neural Resource a Poisson process to describe the system’s stochasticity. While Poisson noise provides a more accurate approximation to neuronal population activity (e.g., Softky & Koch, 1993), it can also be closely approximated by Gaussian noise with an appropriate scaling of variability, and a variant of the Neural Resource model using this approximation has been shown previously to provide a similar account of WM recall errors (Schneegans & Bays, 2017; Taylor & Bays, 2020). Secondly, the two models use different decoding rules, with TCC using the MAX rule (i.e., a “winner-takes-all” decision rule) and Neural Resource using the maximum likelihood decoder. The latter decoder is asymptotically unbiased and minimal in variance, and therefore typically preferred over the generally suboptimal MAX rule (Kay, 1993).

Concepts of similarity and generalization span sensation and perception to a variety of higher cognitive processes. For example, some of the most fundamental perceptual organization principles, such as the Gestalt similarity principle, depend on the feature similarity of perceived objects (Ellis, 1938). Similarly, processes including, but not limited to object recognition (Ullman, 1989), categorization (Hintzman, 1986), and category learning (Gluck, 1991) are often assumed to require judging the similarity of perceptual or semantic representations. Moreover, adequate generalization across stimuli is crucial for adaptive behavior, and it has been shown that psychiatric disorders such as anxiety disorder are characterized by an excessive generalization of fear memories to unrelated contexts (Ghosh & Chattarji, 2015). Although our study found compelling evidence that the psychophysical similarity function does not explain the shape of working memory error distributions, confusability of stored items nonetheless plays a significant role in memory retrieval. Indeed comparison of a visible cue with features in memory is a critical component of analogue report tasks, with failures in this process leading to transposition or “swap” errors. Successful accounts of these errors have been formulated in terms of both feature similarity (Oberauer & Lin, 2017) and population coding (Schneegans & Bays, 2017), and future work could aim to synthesize these approaches.

More generally, an abundance of research has demonstrated shared mechanisms and close interplay between perception and memory. Neuroimaging studies have extensively investigated the extent to which the brain areas involved in sensory processing also play a critical role in the short-term storage of visual information (Serences, 2016; Xu, 2017). Moreover, research has shown that the contents of VWM can alter the perception of stimuli (Kang, Hong, Blake, & Woodman, 2011; Teng & Kravitz, 2019), and that perceptual input can disturb VWM representations (Lorenc, Mallett, & Lewis-Peacock, 2021; Rademaker, Bloem, De Weerd, & Sack, 2015). Several studies have shown that encoding precision and resource allocation in VWM depends on low-level sensory characteristics of stimuli in a similar way to perception (Bays, 2016; Tomić & Bays, 2018).

There is an important conceptual distinction to be made between psychophysical similarity and perceptual variability. The former is described by a psychophysical distance scaling function, which can be estimated from quad, triad or Likert tasks and is approximately exponential with physical distance, reflecting a non-linear internal representation of the distances between features, distinct from non-uniformities in the representation of the features themselves (Appelle, 1972; Bae, Olkkonen, Allred, Wilson, & Flombaum, 2014; Panichello, DePasquale, Pillow, & Buschman, 2019). The latter is measured using perceptual discrimination or analogue report tasks, where non-uniformity is observed as differences in accuracy and precision when different features in a stimulus space are tested.

Non-uniformity in internal representation of feature spaces has been extensively investigated (Ganguli & Simoncelli, 2014; Girshick, Landy, & Simoncelli, 2011; Wei & Stocker, 2015, 2017) and shown to influence both perceptual and WM errors (although it does not provide a sufficient explanation for long tails). An extension of the Neural Resource model incorporating natural image statistics has been shown to quantitatively reproduce non-uniformities in WM (Taylor & Bays, 2018). In contrast, to our knowledge no study prior to Schurgin et al. (2020) has suggested a relationship between the psychophysical similarity function and WM errors. It is this purported relationship, and the claim that non-linearity of the similarity function explains the shape of WM error distributions, that we test here and find evidence against.

A possible source of confusion between these distinct concepts lies in the fact that variation in perception of individual features affects estimates of the psychophysical similarity function obtained from tasks such as the quad task. This influence is thought to be responsible for small deviations in these estimates from the idealized exponential relationship with distance (Nosofsky, 1992). The source study recognised the existence of these deviations, and arguably should have attempted to remove them so the TCC response function could more accurately reflect the underlying psychophysical distance function, however we followed the source study and allowed them to contribute to the response function, where by smoothing the peak of the response function they may actually have made the Empirical-TCC model a better fit to WM data.

Understanding how uncertainty of recall is represented in the brain and used to guide decisions has become an important topic of working memory research (Honig, Ma, & Fougnie, 2020; H.-H. Li, Sprague, Yoo, Ma, & Curtis, 2021). In addition to error distribution, population coding accounts have been shown to make accurate predictions about subjective confidence and its correlation with error (also called meta-cognitive accuracy; Bays, 2016; Schneegans et al., 2020; van den Berg, Yoo, & Ma, 2017). In the case of Poisson noise, the uncertainty associated with a recall estimate can be determined from the total activity on which it is based (i.e. the spike count) or, more generally, from variation in the width of the associated likelihood function (providing a link to variable precision models; Fougnie et al., 2012; van den Berg et al., 2012). In the TCC model, the amplitude of the maximum response has been proposed to play the equivalent role in determining subjective confidence, although the justification for ignoring the amplitude of other responses is unclear, and a comparable quantitative match to empirical data has not yet been demonstrated.

